# Improved Graph-Based Antibody-Aware Epitope Prediction with Protein Language Model-Based Embeddings

**DOI:** 10.1101/2025.02.12.637989

**Authors:** Mansoor Ahmed, Sarwan Ali, Avais Jan, Imdad Ullah Khan, Murray Patterson

## Abstract

The accurate identification of B-cell epitopes is critical in antibody design, diagnostics, and immunotherapies. Many *in silico* approaches have recently been proposed to predict epitopes, but these approaches struggle primarily because of the variational and conformational nature of epitopes. However, deep learning-based approaches have recently shown great promise in achieving better performance at the epitope prediction task. In this paper, we employ a graph convolutional network (GCN) coupled with pre-trained protein language model (PLM)-based embeddings for epitope prediction on a benchmark antibody-specific epitope prediction (AsEP) dataset. We explore the use of different PLM-embedding methods on the epitope prediction task and show that the choice of PLM embeddings impacts the performance. Specifically, we find that antibody-specific PLMs such as AntiBERTy and general PLMs such as ProtTrans and ESM-2 for antigens provide improved epitope prediction performance with an AUCROC of 0.65, precision of 0.28, and recall of 0.46. The source code is available at: https://github.com/mansoor181/walle-pp.git.

## 1 Introduction

Antibodies are large, Y-shaped proteins produced by B-cells that play a critical role in the immune system by identifying and neutralizing foreign substances such as toxins, bacteria, and viruses, collectively known as antigens. Antibodies are currently known to be the largest class of biotherapeutics, where five of the current top 10 blockbuster drugs are monoclonal antibodies [1]. Antibodies also have wide applications in diagnostics and biological research and are traditionally produced *in vivo* by immunizing animals [2,3]. However, these approaches are time-consuming, expensive, and laborious and make it imperative to develop *in silico* methods for antibody design by exploiting recent advances in artificial intelligence, which can potentially reduce the antibody design search space by producing cost-effective antibodies [4,5,6].

A critical step in antibody design is antigen binding site or epitope prediction, which involves identifying the residues on the surface of an antigen that are recognized and bound by an antibody [7]. Accurate epitope prediction is essential for understanding antibody-antigen interactions and designing antibodies with high binding affinity [5]. However, this problem is inherently challenging due to several factors [8,9]. First, epitopes are often non-linear and conformationally diverse, making it difficult to capture their spatial relationships using sequence-based methods alone [6]. Second, a single antigen can have multiple epitopes, which further complicates the prediction task [1]. Finally, most of the current approaches rely on sequence-based data for epitope prediction, which fail to model the complex spatial arrangements of antigen binding sites [2].

Current approaches to epitope prediction can be broadly categorized into sequence-based and structure-based methods [10]. Sequence-based methods rely on amino acid sequences to predict epitopes, while structure-based methods leverage the 3D structure of antigens to identify potential binding sites [11]. Additionally, these methods can be classified as antibody-agnostic or antibody-aware [12]. Antibody-agnostic methods predict epitopes without considering the specific antibody, whereas antibody-aware methods incorporate information about the antibody’s structure and binding interaction [3]. The latter approaches have proved to be more accurate as they enable the identification of specific epitopes on the antigen surface that are recognized by a particular antibody [1]. Graph-based approaches, which represent antigens and antibodies as graphs, have shown further promise due to their ability to model complex spatial relationships [7,10]. However, the choice of embedding methods for encoding residues in these graphs is critical for achieving accurate predictions [13].

In this work, we explore the use of protein language models (PLMs) for embedding residues in a graph-based antibody-aware epitope prediction setup. Specifically, we evaluate the performance of pre-trained PLMs such as ESM-2 [14], AntiBERTy [15], ProtTrans [16], and ESM-IF [17], alongside classical embedding methods like one-hot encoding and BLOSUM62. We extend the WALLE [13] framework by integrating these embeddings into a graph convolutional network (GCN) to predict epitope residues on antigen surfaces. We pose the epitope prediction problem as a link prediction task: for each node in the antibody and antigen graphs, predict whether there is an edge between them or not. The nodes in the antigen graph participating in the binding are classified as epitopes. Our main contributions are as follows:

1. We provide a comprehensive comparison of different PLM-based embedding methods for the epitope prediction problem.
2. We propose the use of AntiBERTy for antibody residue embedding and ProtTrans for antigen embedding, along with the GCN model for improved epitope prediction.

The remainder of this paper is organized as follows. Section 2 discusses related literature on epitope prediction and graph-based methods. Section 3 presents the proposed approach with the problem formulation, data representation, and model architecture. Section 4 discusses the dataset and model training details. Section 5 discusses the experimental results and comparisons with baseline methods and outlines future research directions. Finally, Section 6 concludes the paper.

## 2 Related Work

Antibody-agnostic methods often rely on sequential, structural, or a combination of both features but lack the specificity required for antibody-specific applications [11]. For instance, *epitope1D* [3], *GraphBepi* [7], and *EpiGraph* [10] are prominent B-cell epitope prediction methods using graph-based representations that do not incorporate antibody-specific information. On the other hand, antibody-aware methods such as *EpiScan* [8], *PECAN* [18], and *EPMP* [11] explicitly incorporate information about the antibody’s structure or sequence to predict epitopes.

Graph-based approaches have emerged as a powerful paradigm for epitope prediction, capturing the inherent spatial and sequential relationships in protein structures. Methods like *PECAN* [18], *PInet* [19], and [20] uses graph neural networks (GNNs) to predict protein-protein interaction interfaces, including antibody-antigen binding sites by employing attention mechanisms. *EPMP* [11] is a neural message-passing framework that uses asymmetrical architectures for paratope-epitope prediction, while *EpiPred* [12] combines conformational matching and antibody-antigen scoring to improve epitope prediction results. Moreover, [21] uses GNN to capture information on spatial neighbors of a target residue and attention-based didirectional long shortterm memory (Att-BLSTM) networks to extract global information from the whole antigen sequence.

Some recent approaches have combined PLM-based embeddings and graph-based architectures for epitope prediction. *EpiGraph* [10] uses graph attention networks (GAT) with ESM-2 and ESM-IF embeddings to predict epitopes, achieving state-of-the-art performance. Similarly, [13] introduced *AsEP*, a benchmark dataset and a graph-based method (*WALLE*) that employs embeddings from ESM-2 and AntiBERTy in a GNN setup for antibody-specific epitope prediction. *GraphBepi* [7] is another graph-based model that leveraged ESM-2 embeddings for epitope prediction. *DeepProSite* [9] uses *ESMFold* [14] for protein structure prediction and employs embeddings from pre-trained ProtTrans to predict protein binding sites using graph transformers, while ESMBind [22] finetunes complex ESM-2 models ranging from 8M to 650 parameters on an annotated protein binding sites dataset for protein binding site prediction task.

## 3 Proposed Approach

This section presents the problem formulation of the epitope prediction task, the data representation strategies employed, and the overall architecture of the proposed method.

### 3.1 Problem formulation

The interaction between antibodies and antigens can be framed as predicting links between the vertices of two disjoint undirected graphs: an antibody graph *G*_*A*_ = (*V*_*A*_, *E*_*A*_) and an antigen graph *G*_*B*_ = (*V*_*B*_, *E*_*B*_), *i*.*e*., *V*_*A*_ ∩ *V*_*B*_ = ∅. Here, *V*_*A*_ and *V*_*B*_ represent the sets of vertices (residues) for the antibody and antigen graphs, respectively, while *E*_*A*_ and *E*_*B*_ represent their sets of edges based on residue proximity. Each vertex is encoded into a vector using a function 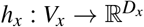 where *x* ∈ {*A, B*} and *h* can be any encoding function such as one-hot encoding or pre-trained embeddings from a PLM. The adjacency matrix 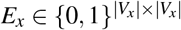 is derived from the distance matrix of the residues of antigen or antibody. Each entry *e*_*i j*_ indicates the proximity between residue *i* and residue *j*; *e*_*i j*_ = 1 if the Euclidean distance between any non-hydrogen atoms of residue *i* and residue *j* is less than 4.5Å, and *e*_*i j*_ = 0 otherwise. In order to make the computations faster, the antibody graph *G*_*A*_ is constructed only from the CDR residues of the antibody’s heavy and light chains, while the antigen graph *G*_*B*_ is constructed from the surface residues of the antigen. We address the following two key tasks in this setup:

#### Epitope Node Prediction

This task involves identifying antigen residues that interact with the antibody. A residue *v* ∈ *V*_*B*_ is labeled as an epitope (1) if it is within 4.5Å of any residue in *V*_*A*_; otherwise, it is labeled as a non-epitope (0). Formally, the task is a binary node classification problem on the antigen graph *G*_*B*_, conditioned on the antibody graph *G*_*A*_. The classifier *f* : *V*_*B*_ → {0, 1} is defined as:

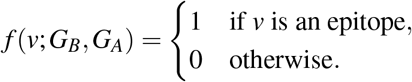

#### Link Prediction

This task predicts interactions between antibody and antigen residues, forming a bipartite graph *K*_*m,n*_ = (*V*_*A*_,*V*_*B*_, *E*), where *m* = |*V*_*A*_|, *n* = |*V*_*B*_|, and *E* represents potential inter-graph edges. An edge *e* ∈ *E* is labeled as 1 if the corresponding residues are within 4.5Å of each other and 0 otherwise. The task is formulated as a binary edge classification problem, with the classifier *g* : *K*_*m,n*_ → {0, 1} defined as:

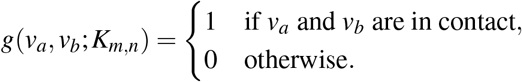

The problem formulation is derived partially from [10,13,18].

### 3.2 Graph Representation

Each antibody-antigen complex is represented as a graph, where protein residues are modeled as vertices. Edges are drawn between residues if their non-hydrogen atoms are within 4.5Å of each other. We focus on surface residues, excluding buried residues with zero solvent-accessible surface area from the antigen graph, and non-CDR residues from the antibody graph.

In our study, we employ various embedding methods to represent protein residues as continuous vectors. These methods include both classical techniques and state-of-theart PLMs. Specifically, we use AntiBERTy^4^, an antibody-specific transformer language model pre-trained on 558M natural antibody sequences, which produces embeddings of fixed size *L* × 512 where *L* represents the length of the sequence. Additionally, we utilize ESM-2^5^, a PLM that captures the evolutionary features of the protein by generating embeddings of size *L* × 1280 using the `ESM-2_t33_650M_UR50D` model. Conversely, WALLE [13] used a smaller ESM-2 model `ESM-2_t12_35M_UR50D` to produce 480-dimensional sequence embeddings. We also employ ESM-IF1, an inverse folding model, to produce 512-dimensional structure embedding vectors with the model `esm_if1_gvp4_t16_142M_UR50`. ProtTrans^6^ is a self-supervised transformer-based autoencoder fine-tuned on more than 500 million protein sequences and is used to generate 1024-dimensional embeddings with the model `prot_t5_xl_uniref50`. For comparison, we also include classical sequence embedding methods such as BLO-SUM62 and One-Hot Encoding.

### 3.3 Model Architecture

The overall architecture of our graph-based framework for antibody-aware epitope prediction is shown in Figure 1. The model takes as input an antibody-antigen graph pair, constructed as detailed in Section 4.1, and performs node- and edge-level predictions.

**Fig. 1:**
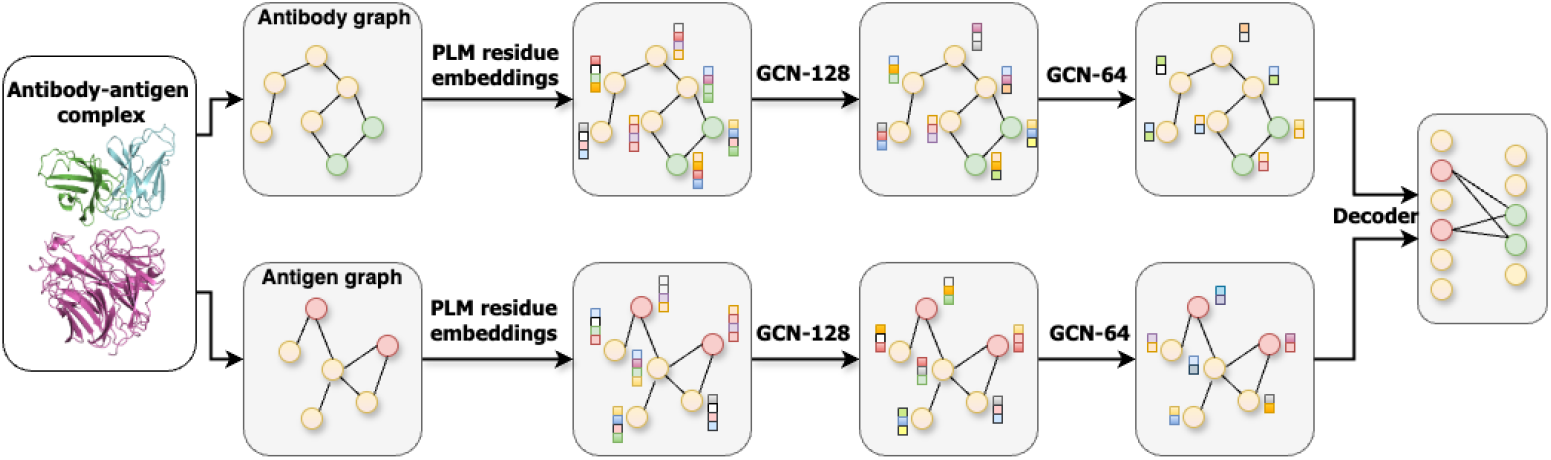
The schematic of representing an antibody-antigen complex structure as a graph pair, employing pre-trained protein language model (PLM)-based embeddings and the graph convolutional network (GCN) model architecture. The yellow-colored nodes represent non-binding residues, while green and red-colored nodes represent paratopes and epitopes, respectively.

We incorporate graph modules that process the input graphs of antibody and antigen structures separately, inspired by WALLE [13], PECAN [18], and EPMP [11]. The antibody graph is represented by node embeddings 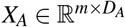 and an adjacency matrix *E*_*A*_, while the antigen graph is described by node embeddings 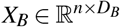 and its corresponding adjacency matrix *E*_*B*_. The embedding vector sizes *D*_*A*_ and *D*_*B*_ vary depending on the embedding method used, such as AntiBERTy, ESM-2, or other methods.

Both antibody and antigen graph nodes are first projected into a dimensionality of 128 using fully connected layers. The resulting embeddings are then passed through two GCN modules consecutively to refine the features and yield updated node embeddings 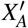 and 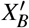 with a reduced dimensionality of *m* × 64 and *n* × 64, respectively. The output from the first GCN layer is passed through a ReLU activation function while outputs from the second GCN layer are directly fed into the decoder module. These GCNs operate independently, each with its own parameters, ensuring that the learned representations are specific to the antibody or the antigen. The use of separate GCN modules for the antibody and antigen allows for the capture of unique structural and functional characteristics pertinent to each molecule before any interaction analysis. This design choice aligns with the understanding that the antibody and antigen have distinct roles in their interactions, and their molecular features should be processed separately.

A decoder is employed to predict the binary labels of edges between the antibody and antigen graphs. The decoder takes a pair of node embeddings output by the graph modules, 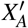 and 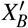, as input and constructs a bipartite adjacency matrix of size *m* × *n*. Then, it predicts the probability of each edge: an edge is assigned a binary label of 1 if the predicted probability is greater than 0.5 or 0 otherwise. For the epitope prediction task, edge-level predictions are converted to node-level by summing the predicted probabilities of all edges connected to an antigen node. An antigen node is assigned a label of 1 if the number of connected edges is greater than a threshold or 0 otherwise. The threshold is treated as a hyperparameter and is optimized in the experiments.

## 4 Experimental Setup

In this section, we provide the dataset description, accompanied by an exploratory data analysis and dataset split strategies, and the implementation details of our framework.

### 4.1 Dataset

We utilized the AsEP dataset [13], a novel benchmark dataset of antibody-antigen complexes designed specifically for epitope prediction tasks. The dataset was derived from the Antibody Database (AbDb) and the Protein Data Bank (PDB), comprising 1,723 unique antibody-antigen complexes after filtering and deduplication. Each complex includes a single-chain protein antigen paired with a conventional antibody containing both heavy and light chains. Non-canonical residues and unresolved CDR regions were excluded to ensure data quality.

#### Exploratory Data Analysis (EDA)

Our EDA revealed several key insights into the dataset and are shown in Figure 2. The distribution of epitope residues showed a mean of 19 ± 4.7, while the antigen surface residues numbered in the hundreds. The contact distribution between residues in the bipartite graph had a mean of 43.7 contacts with a standard deviation of 12.8. Additionally, the dataset includes 641 unique antigens and 973 epitope groups, highlighting the diversity and complexity of the antibody-antigen interactions captured in the AsEP dataset.

**Fig. 2:**
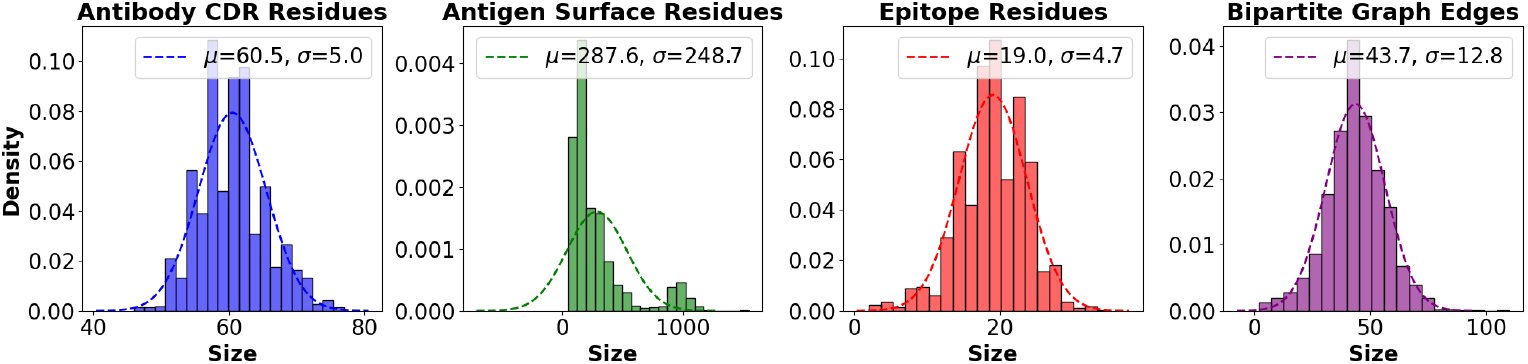
(left to right) The distribution of the size of antibody CDR residues, antigen surface residues, epitope residues, and bipartite graph edges (antibody-antigen residueresidue contacts) in the dataset. The dotted lines represent the fitted normal distribution.

#### Dataset Split

We employed two dataset splits for the model training and evaluation:

1. **Epitope to Antigen Surface Ratio:** This split ensures a similar distribution of epitope to non-epitope nodes across the training, validation, and test sets, with 1383 complexes for training and 170 complexes each for validation and testing.
2. **Epitope Groups:** This split challenges the model to generalize to unseen epitope groups, with 641 unique antigens and 973 epitope groups having multi-epitope antigens, each treated as an antigen-antibody pair. The dataset was divided into training, validation, and test sets with an 80/10/10 split, resulting in 1383 complexes for training and 170 complexes each for validation and testing.

### 4.2 Implementation Details

The model implementation details were followed in part from WALLE [13]. We used the PyTorch Geometric framework to build the graph modules in the proposed model. We used a weighted loss function consisting of a sum of binary cross-entropy loss for the bipartite graph link reconstruction and a regularizer for the number of positive edges in the reconstructed bipartite graph to train the model. We used the mean of edges in the antibody-antigen bipartite graph (as shown in Figure 2) to set the regularizer in the loss function to constrain the model not to predict too many or too few positive edges. We used two configurations for the decoder: a fully connected linear layer (GCN-L) and an inner product decoder (GCN). We used the same set of hyperparameters and loss functions for both dataset split settings. We evaluated the performance of our model using the standard classification metrics of Matthews correlation coefficient (MCC), the area under the precision-recall (AUPRC) curve, the area under the ROC curve (AUCROC), precision, recall, F1 score, and balanced accuracy (BACC). All experiments were run in parallel on a 40-CPU research server with Ubuntu 16.04, and it took an average of 1.5 hours for embedding generation and 50 minutes for training on one configuration.

## 5 Results and Discussion

In this section, we report and discuss the results of our experiments on epitope node classification and bipartite link prediction tasks under epitope-to-antigen surface ratio and epitope group split settings using standard classification metrics.

### 5.1 Epitope Node Prediction

For epitope node classification, PLM embeddings demonstrated superior performance compared to classical encoding methods across both split strategies. Under epitope ratio dataset split (Table 1-group a), AntiBERTy/ProtTrans embeddings configuration for an antibody-antigen pair achieved the highest MCC of 0.263 and AUCROC of 0.65, outperforming other embedding configurations. These results were consistent for both configurations of the decoder in our model, *i*.*e*., GCN and GCN with a fully connected linear layer. AntiBERTy consistently provided improved performance for antibody embeddings on the metrics of MCC, AUPRC, precision, and F1 score, while ProtTrans showed better performance for antigen embeddings across different combinations on these metrics. These results are consistent with [9,23], suggesting that ProtTrans embeddings provide a rich context for binding site prediction. It was also observed that ESM-IF embeddings for antigens demonstrated very poor performance on all metrics except recall, while it provided average performance in the case of antibodies.

**Table 1:**
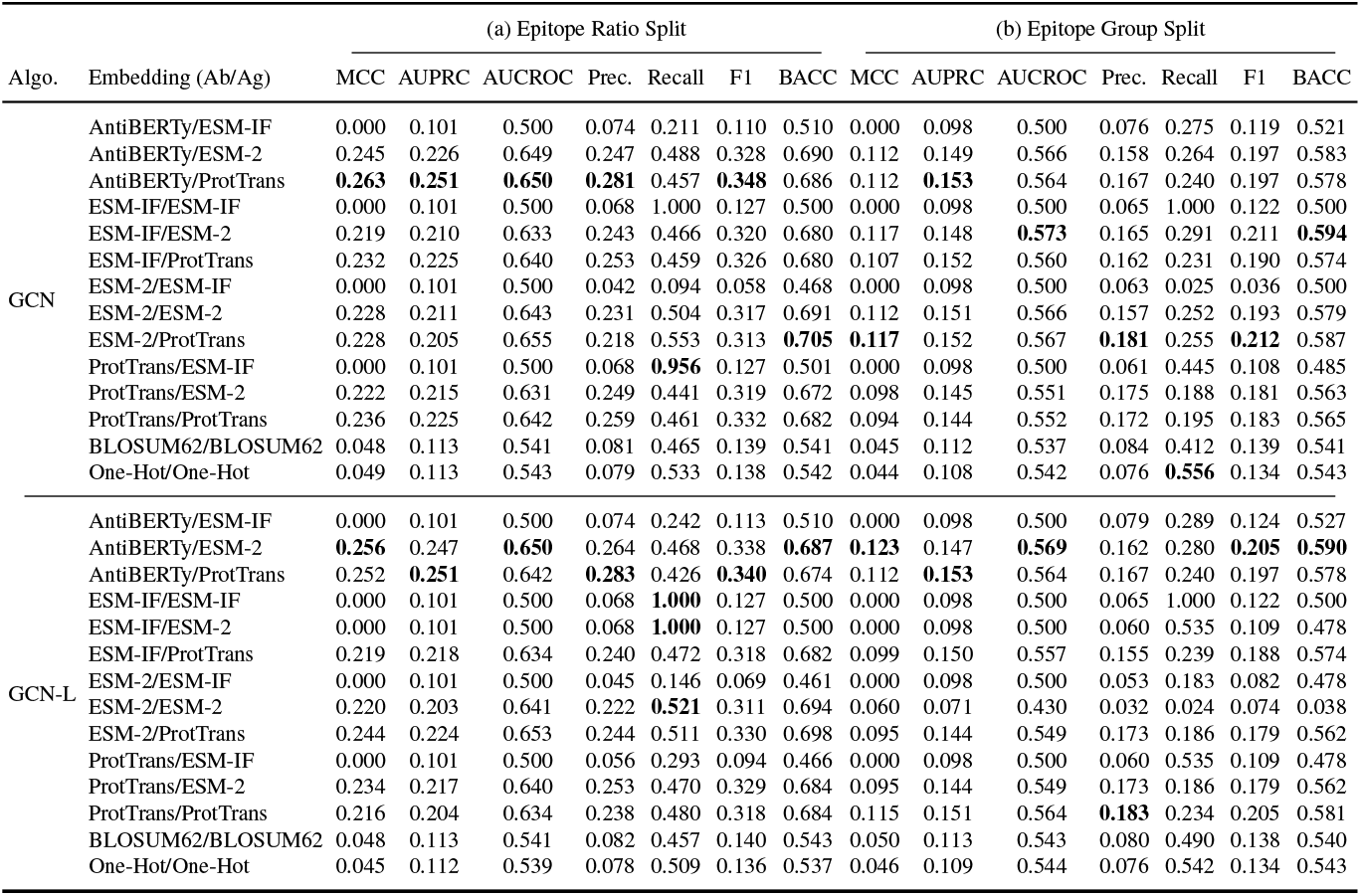
Performance of our model on the test set for epitope node prediction with epitope to antigen surface ratio (epitope ratio) and epitope group split strategies. The best values are highlighted in bold.

The more challenging epitope group-based split (Table 1-group b) revealed interesting tradeoffs. The results were not consistent for a specific configuration of embeddings on the majority of evaluation metrics. In general, ESM-2/ProtTrans led to the best results on the metrics of MCC, AUPRC, precision, F1 score, and balanced accuracy (BACC). Similar to the observations in the case of the epitope-ratio split, employing AntiBERTy for antibodies consistently led to better overall results on all metrics in the epitope group split. Consistent with the observations in [13], we also observed that the model struggled with the epitope group split setting compared to the epitope ratio because it could not generalize well on unseen epitopes and showed biases towards the training set. While our model with ESM-2 embeddings attained the best results overall on these metrics, classical one-hot encoding surprisingly achieved the highest recall (0.556), suggesting simpler methods may capture broader structural patterns beneficial for unseen epitope detection. This dichotomy highlights the need for hybrid approaches combining PLM-derived semantics with geometric features [8,19].

### 5.2 Link Prediction

The link prediction results (Table 2) demonstrated the task’s inherent complexity, with maximum MCC scores of 0.126 and 0.054 for ratio and group splits, respectively. The performance gap between tasks suggests node classification benefits more from local residue features, while link prediction requires finer-grained interaction patterns that current GCN architectures struggle to capture. Consistent with the epitope prediction task, the overall results for bipartite link prediction demonstrated that the AntiB-ERTy/ProtTrans embedding configuration achieves the best performance across almost all metrics for epitope-ratio split. Contrary to epitope node classification, we observed very high AUCROC scores of 0.852 for bipartite link prediction, where we were able to get a maximum AUCROC of 0.65. It was also observed that classical embedding methods such as BLOSUM62 and one-hot encoding showed degraded performance for both dataset splits and GCN decoder configurations on all metrics except AUCROC. Overall, these results suggest that GCN modules with a simple decoder performing the inner product of antibody and antigen embeddings provide consistent and best results compared to GCN with a linear layer as the decoder for both epitope node prediction and bipartite link prediction tasks. Consistent with the investigations in [10,13], our results demonstrated that the choice of the PLM embedding method impacts the model performance on the epitope prediction and bipartite link prediction tasks.

**Table 2:**
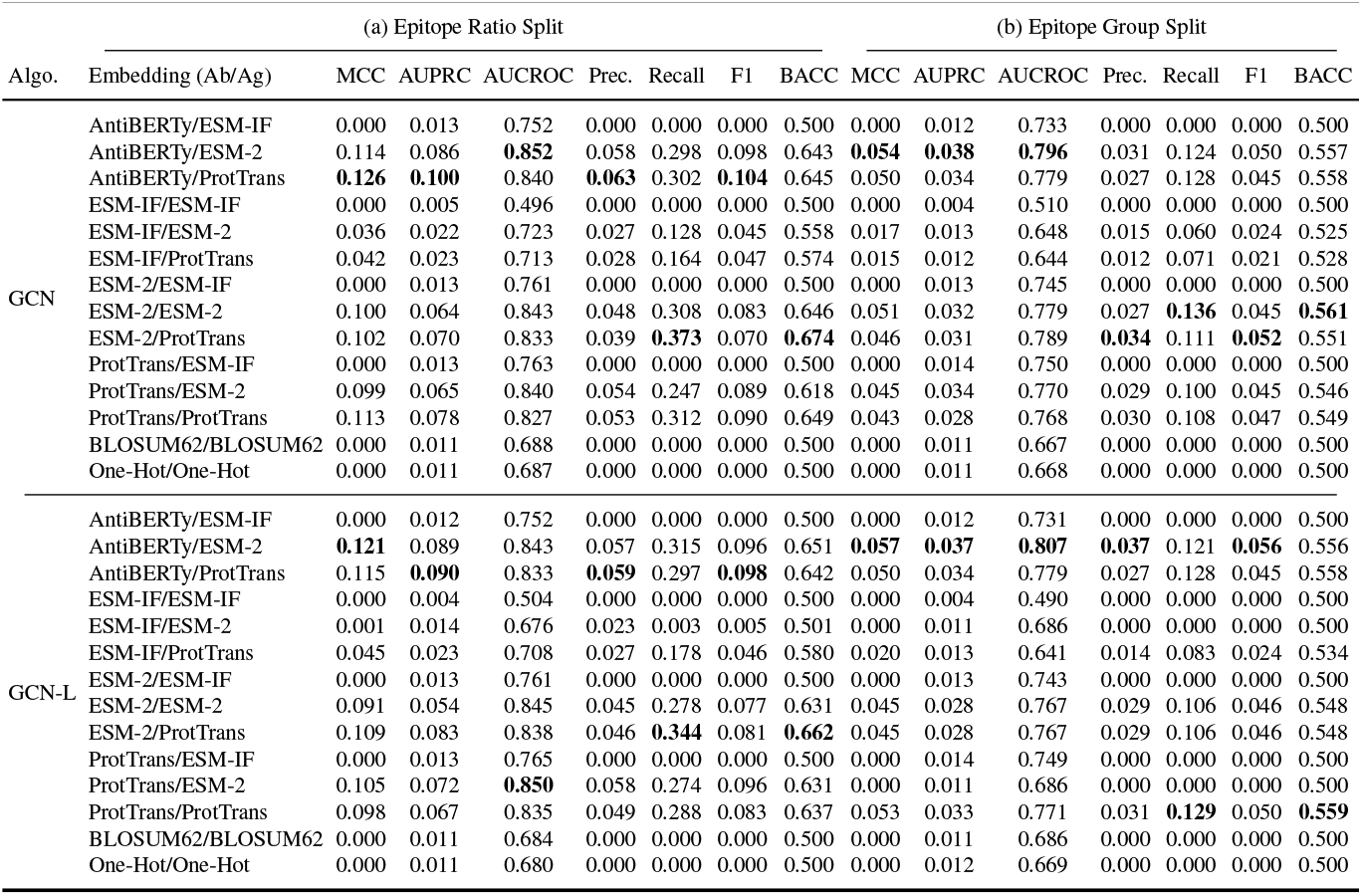
Performance of our proposed model on bipartite link prediction task. The best values are highlighted in bold.

### 5.3 Baselines Comparison

We compared our proposed approach with established baselines (WALLE^7^ [13], EpiPred [24], MaSIT-site [25], and ESMBind [22]) and revealed significant performance improvements (Table 3). Readers are referred to the related work (Section 2) for further description of these different baseline methods. For the epitope ratio split, our proposed approach achieved an MCC of 0.263, outperforming WALLE [13] and the next best model, EpiPred [12]. Notably, our model’s superior performance extended across all metrics, with a precision of 0.281 and recall of 0.457, indicating a balanced ability to identify true epitopes while minimizing false positives. The AUC-ROC of 0.65 further confirmed our approach’s robust epitope predictive capability, significantly surpassing ESMBind’s 0.506. PECAN [18] originally employed a GNN architecture for epitope prediction and achieved a precision of 0.154 without PLM embeddings. This improvement suggests that the integration of GCN with different PLMs captures critical structural and sequential information about antigens and antibodies that were previously not explored in detail. Moreover, the performance gap between our proposed model and sequence-based methods like ESMBind (MCC of 0.016) underscores the importance of incorporating structural information for accurate epitope prediction [7,10].

**Table 3:**
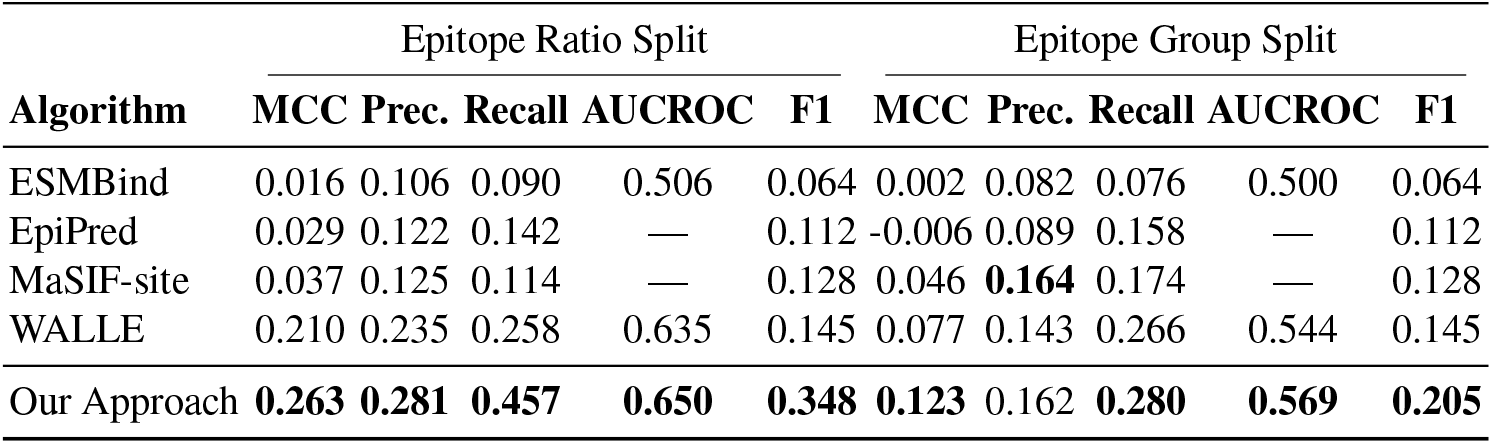
Performance comparison of our proposed model with different baselines on two dataset split strategies. The values in bold represent the best values, while the dash represents that these metrics were not computed.

Though the performance dropped in the challenging epitope groups split across all methods, many of which struggled to achieve positive MCC values, our model maintained its lead with an MCC of 0.077. MaSIF-site [25] showed competitive performance, suggesting that its geometric deep learning approach captures some generalizable epitope features. This suggests that current approaches, including WALLE, may be overfitting to epitope patterns seen in training data, emphasizing the need for more diverse training sets and potentially more sophisticated regularization techniques.

### 5.4 Limitations and Future Research Directions

Though our results demonstrated performance improvement for epitope prediction, there are limitations to our approach, and hence we note some potential future research directions. The authors in [10] demonstrate that the concatenation of ESM-2 and ESM-IF1 embeddings improved the epitope prediction performance. In the future, we will explore the integration of different PLM-based embedding methods, such as ESM-2 with ProtTrans, for antibody-aware epitope prediction using separate GAT modules. Other deep learning models, such as bi-directional long short-term memory (BiLSTM [7]), have shown potential for predicting binding sites and need to be explored with concatenated PLM embeddings for the epitope prediction task. There is also immense potential in exploring other geometric deep learning models, such as diffusion-convolutional neural networks (DCNN [26]) for epitope prediction. We will also explore other protein representation approaches, such as point clouds and meshes [25], that have demonstrated success for protein binding site prediction [19].

## 6 Conclusion

We extended a graph-based approach (WALLE) for antibody-aware epitope prediction on a novel benchmark dataset by exploring different pre-trained PLMs and classical embedding methods. We show that the use of PLM-based embeddings integrated with the graph-based models provides improved epitope prediction performance. Specifically, our experiments with different state-of-the-art PLM-based embedding methods show that using AntiBERTy for antibody embedding and ProtTrans for antigen embedding improves the epitope classification accuracy. We also discuss potential future research directions for improved epitope prediction.

## Footnotes

4 https://github.com/jeffreyruffolo/AntiBERTy.git

5 https://github.com/facebookresearch/esm.git

6 https://github.com/agemagician/ProtTrans.git

7 The code for the preprint version https://arxiv.org/abs/2407.18184v1 was the latest available for comparison at the time of publication.

